# Whokaryote: distinguishing eukaryotic and prokaryotic contigs in metagenomes based on gene structure

**DOI:** 10.1101/2021.11.15.468626

**Authors:** Lotte J. U. Pronk, Marnix H. Medema

## Abstract

Metagenomics has become a prominent technology to study the functional potential of all organisms in a microbial community. Most studies focus on the bacterial content of these communities, while ignoring eukaryotic microbes. Indeed, many metagenomics analysis pipelines silently assume that all contigs in a metagenome are prokaryotic. However, because of marked differences in gene structure, prokaryotic gene prediction tools fail to accurately predict eukaryotic genes. Here, we developed a classifier that distinguishes eukaryotic from prokaryotic contigs based on foundational differences between these taxa in gene structure. We first developed a random forest classifier that uses intergenic distance, gene density and gene length as the most important features. We show that, with an estimated accuracy of 97%, this classifier with principled features grounded in biology can perform almost as well as the classifiers EukRep and Tiara, which use k-mer frequencies as features. By re-training our classifier with Tiara predictions as additional feature, weaknesses of both types of classifiers are compensated; the result is an enhanced classifier that outperforms all individual classifiers, with an F1-score of 1.00 on precision, recall and accuracy for both eukaryotes and prokaryotes, while still being fast. In a reanalysis of metagenome data from a disease-suppressive plant endosphere microbial community, we show how using Whokaryote to select contigs for eukaryotic gene prediction facilitates the discovery of several biosynthetic gene clusters that were missed in the original study. Our enhanced classifier, which we call ‘Whokaryote’, is wrapped in an easily installable package and is freely available from https://git.wageningenur.nl/lotte.pronk/whokaryote.

## Introduction

Microbiomes are increasingly recognized for playing a large role in the health and development of their hosts^1,2^. Studies aiming to characterize these microbial communities are more and more shifting from largely marker-gene-based community abundance profiling (e.g., 16S, ITS) to analyses of the complete metagenome using shotgun sequencing. Analyzing the complete metagenome has provided valuable insights in microbial ecology and microbiome-host interactions^3–5^.

Many metagenomic analysis pipelines assume that all contigs will be prokaryotic, while eukaryotic microorganisms are also present in microbiomes^6–9^. Including eukaryotes in metagenome analyses could provide a more complete picture of their ecological role and functional capabilities. To study eukaryotic sequences in metagenomes, a reliable taxonomic classification method on the contig level is needed.

Most current metagenome taxonomic classification tools were designed mostly with prokaryotes in mind. Tools such as CAT/BAT^10^, DIAMOND+MEGAN^11^, or Kraken2^12^ are good at assigning taxonomy up to the species level to metagenomic contigs, but they require sequence homology searches (CAT, DIAMOND+MEGAN) or use k-mers (Kraken2) and thus require the use of large pre-existing databases. These tools are therefore relatively time-consuming to run on large metagenomic datasets, and uncultivated organisms are frequently missed because they are not in any database yet. Additionally, some sequence-homology-based methods, like CAT, require accurate gene predictions, which paradoxically requires knowledge on at least the empire (eukaryote/prokaryote) level for each contig to select the best gene predictor. Prokaryotic gene predictors do not consider introns, making them unsuited for predicting eukaryotic genes. The eukaryotic gene predictor Augustus^13^ uses models that were trained on specific organisms and can be trained on metagenomic bins to predict the genes in such a bin, an approach used in the EukRep pipeline^14^. However, there is no good solution for de novo gene prediction in unbinned contigs. MetaEuk predicts protein-coding genes on eukaryotic metagenomic contigs, but relies on a large and likely incomplete reference database of protein profiles^15^. Because of the limitations of current methods, many metagenomic studies may be missing eukaryotic genes altogether because of wrongly annotating them. K-mer based methods such as Kraken2 do not rely on gene predictions, but they require a database of closely related reference genomes. There are not many reference genomes of eukaryotic microorganisms available in public databases yet, which could lead to an underestimation of eukaryotes in microbiomes.

In order to allow metagenomic analysis pipelines to selectively run eukaryote-specific gene finding on all contigs likely to be eukaryotic (whether belonging to a known taxon or not), simply classifying metagenomic contigs as either prokaryotic or eukaryotic (instead of comprehensively assigning taxonomy) would be a logical solution. Some efforts have recently been made in this direction. For example, EukDetect maps reads to a database of universal eukaryotic marker genes to detect eukaryotes in metagenomes^16^, but this approach will not find contigs that do not contain these marker genes. The tool EukRep^14^ calculates k-mer counts of 5-kb fragments of (meta)genomic contigs and then classifies the complete contig based on a majority vote using a support vector machine model. The authors show that using eukaryotic gene predictors on sequences classified as eukaryotic leads to more accurate annotations and downstream analyses, including higher-level taxonomic classification^14^. Despite its high overall accuracy, EukRep does not perform very well on several prokaryotic and eukaryotic taxa, especially parasites and symbionts. Moreover, it was trained and tested on relatively long contigs, while metagenome assemblies often consist of mostly short contigs. More recently, Tiara, a classifier that uses k-mer counts as features for deep learning models was published^17^, which was shown to be slightly more accurate than EukRep. Tiara was developed with detecting organelle sequences in mind, and therefore has a more fine-grained classification, distinguishing archaea, bacteria, organelles, and eukaryotes. Like EukRep, the performance of Tiara is less accurate on certain organisms. Because the approaches used by both tools do not allow for inference of feature importance, it is very difficult, if not impossible, to determine which k-mers are important for distinguishing between eukaryotes and prokaryotes, and what underlying biological features these k-mers represent. This makes it difficult to correct for any biases. Moreover, k-mer-based approaches by their nature likely require training data from closely related taxa to work well, making it probable that contigs from uncultivated taxa unique to specific biomes will be misclassified.

Here, we introduce a random forest classifier with comparable performance, but which uses manually selected features based on fundamental differences in gene structure between eukaryotes and prokaryotes. The classifier uses intergenic distance, gene density and other genomic features to predict whether a given metagenomic contig belongs to a eukaryote or a prokaryote. Despite the different approach, our tool performs as well and, in some cases, better than EukRep and Tiara on contigs with sizes often found in metagenomes. We also show that using eukaryote-specific tools to analyze the classified eukaryotic contigs can lead to the discovery of microbial traits that have remained undiscovered when solely using prokaryote-oriented tools. Finally, by using Tiara predictions as a feature, we constructed a highly accurate classifier with an F-1 score of > 99.9% on a simulated metagenomic dataset. With this enhanced classifier, which we term ‘Whokaryote’, sequences can be reliably classified for a very wide range of eukaryotes and prokaryotes.

## Methods and Implementation

### Training dataset for the random forest classifier

To train and test the random forest classifier, we downloaded 73 prokaryotic (68 bacteria and 5 archaea) and 25 eukaryotic (12 fungi, 10 protists, 1 plant and 2 animals) genome assemblies from the NCBI GenBank database, see Supplementary Data 1. All non-genomic DNA sequences were removed using the following search terms: API, MIT, mitochondrial, plastid, chloroplast, mitochondrion, non-nuclear, organelle, apicoplast. Next, these genomes were split into non-overlapping artificial contigs with a random length between 5000 and 100000 base pairs according to a triangular distribution with the lower left limit at 5000, the mode at 10000 and the upper right limit at 100000 bp. For the training set, a maximum of 500 contigs per genome were used to prevent an overrepresentation of eukaryotic contigs, which tend to have very large genomes compared to prokaryotes. For every contig (both prokaryotic and eukaryotic), genes were predicted using the prokaryotic gene prediction tool Prodigal^18^ [version] using the metagenomic setting (--meta). We used Prodigal because it is commonly used in metagenomics pipelines already and preparing these output files as input to our classifier will save time in the subsequent gene prediction step. Contigs with one predicted gene or no predicted genes were excluded in further steps (see below for explanation). This resulted in a dataset with a total of 11,285 eukaryotic contigs and 8,403 prokaryotic contigs that we used to train and validate the random forest classifier.

### Used features

Genes in prokaryotes are often packed closely together on the genome in co-regulated operons that are transcribed to a single polycistronic mRNA. Therefore, we expected that genes on eukaryotic contigs generally have a higher intergenic distance and a lower gene density than prokaryotic contigs. Additionally, because genes in operons are under the control of a single promoter, we expected that adjacent pairs of bacterial genes on a given stretch of the genome have a higher probability of being present on the same strand, i.e., they have the same orientation. Because calculation of our selected features requires the presence of at least two genes, contigs with only one predicted gene or no predicted genes are excluded from our classification.

For each contig, the intergenic distance of each gene pair was calculated (start position gene 2 - end position gene 1 = intergenic distance). Next, the mean, standard deviation, and the first and third quartile of the intergenic distance per contig were calculated and used as features for the classifier. The length of every gene was calculated by subtracting the stop position from the start position. The gene density was calculated by dividing the sum of the length of every gene on a contig by the total length of the contig.

In the prodigal gene location output file, genes that are on the positive strand are annotated with a “+” sign, and the genes on the negative strand are annotated with the “-” sign. For every contig, the ratio of genes that are on the same strand was calculated as follows: We determined for every adjacent pair of genes if they are located on the same strand (+ + or - -) or not. The number of pairs that were located on the same strand was divided by the total number of gene pairs. The outcome was used as a feature for the classifier. Finally, the mean gene length of every contig was calculated and used as a feature.

In our enhanced classifier, we used Tiara predictions as an additional feature for training and testing. To obtain Tiara predictions, we used Tiara (version 1.0.2), using the DNA sequence of the contigs as input, and setting the minimum contig length to 3000. We converted them into usable features by converting the predictions to a number: 0 for contigs classified as ‘eukarya’; 1 for contigs classified as ‘prokarya, ‘bacteria’ and ‘archaea’; 2 for contigs classified as ‘unknown’ or ‘organelle’.

Features were also calculated for the reference genome annotations downloaded from NCBI and mapped to the corresponding artificial contigs so that the values could be compared to the prodigal annotations.

### Random forest classifier

Two different classifiers were trained, one without Tiara predictions as a feature and one with Tiara predictions as a feature. All features were stored in a dataframe with contigs as rows and features as columns. To build the random forest classifier, we used the Python package scikit-learn^19^, version 0.23.2. The dataset was randomly split into a training set and a validation set (ratio 70/30), and labels were stratified to maintain the proportion of class labels in the original dataset. Each contig was labeled as ‘eukaryotic’ or ‘prokaryotic’. A random forest with 100 estimators was initialized and 5-fold cross-validation on the training set was used to estimate the expected accuracy. The final model was trained on the training set and tested on the validation dataset. A report of the performance metrics was made using the *metrics*.*classification_report* function of scikit-learn, and feature importance was calculated using the *feature_importances_* function.

### Code availability

The standard random forest classifier and the enhanced classifier are made available in a downloadable Python package that can be used on the Linux command line. The package and instructions to install it are available at https://git.wageningenur.nl/lotte.pronk/whokaryote.

## Results

To train our classifier, we used a dataset of 73 prokaryotic and 25 eukaryotic reference genomes that were fragmented into shorter artificial contigs to resemble a metagenome assembly. Prodigal (metagenomic mode) was used to predict genes on these contigs.

Per contig, we calculated the intergenic distance (average, first quartile, third quartile and standard deviation), gene length (average), gene density, gene length and the percentage of gene pairs with the same orientation. As expected, the intergenic distance was lower on bacterial contigs than on eukaryotic contigs (Figure 1a-d). The gene density was higher on prokaryotic contigs compared to eukaryotic contigs (Figure 1e) and is likely linked to the intergenic distance. The gene length was higher for prokaryotes (Figure 1f). This might be viewed as unexpected, since eukaryotes have introns that can make up a large part of the total gene length; however, we used Prodigal to predict genes on the contigs, which does not consider introns and may predict a gene for every exon, often further reduced in length because the first start codon of an exon may be considerably downstream of the intron/exon junction. Lastly, the percentage of gene pairs on a contig that are in the same orientation was only slightly higher in prokaryotes (Figure 1g), but we still used it for training our classifier.

**Figure 1.**
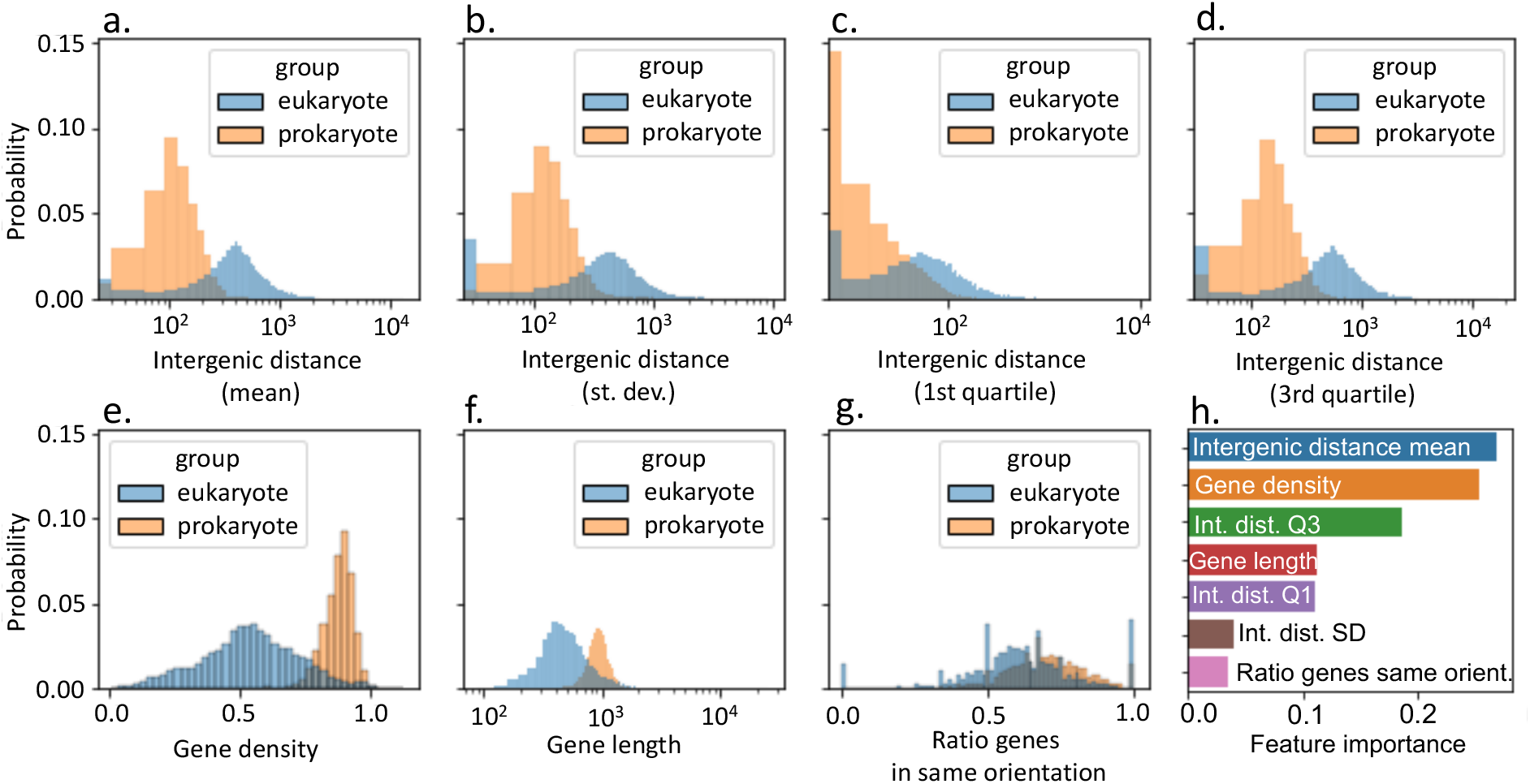
Overview statistics of the features calculated for the training and validation dataset containing gene-based features of artificial contigs taken from 73 prokaryotic and 25 eukaryotic reference genomes. Gene prediction was done using Prodigal-M. The probability distribution of the feature values of all contigs are shown, grouped by the taxonomic group (eukaryote or prokaryote) the contig belongs to. For every contig, the mean, standard deviation, first quartile and third quartile of the intergenic distances between the genes was calculated (a-d, respectively). e: The mean gene density of every contig, calculated by dividing the sum of the length (in base pairs) of all genes by the total length of the contig. f: The mean gene length of every contig, calculated as the end position of the gene minus the start position. g: The ratio of genes on a contig that are on the same strand (e.g., they have the same ‘orientation’). h: The importance of each feature that was used in our random forest classifier trained on artificial contigs of 99 different organisms.

The features we calculated are based on non-ideal gene predictions for eukaryotes, and we wanted to know if the values for each feature we observed are representative and can be used to make claims about biological differences. Therefore, we also calculated the same features for the reference annotations that we mapped to the artificial contigs. Indeed, all features, except for gene length, show (almost) the same distributions in the reference annotations for both eukaryotes and prokaryotes as in the features we calculated with the Prodigal annotations (Supplementary Figure 1). According to the reference annotations, eukaryotes have longer genes than prokaryotes, as is expected because of their introns.

After calculating the features, we trained a random forest with 100 estimators on 70% of the contigs in the training dataset. With the remaining 30%, we tested its performance with a five-fold cross validation, resulting in an average accuracy of 96%. We used this model as our final model. The precision for eukaryotes was 0.97 and for prokaryotes 0.95. Recall was 0.96 for both classes (Table 1). The average intergenic distance, gene density and the 3^rd^ quartile of the intergenic distance were the most important features (Figure 1h).

**Table 1.** Accuracy metrics of contig_classifier, EukRep, Tiara and Whokaryote on a test dataset of artificial contigs taken from the reference genomes of 10 eukaryotes and 15 prokaryotes. P = precision, R = recall, F1 = F1-score.

|  | Contig_classifier |  |  | EukRep |  |  | Tiara |  |  | Whokaryote |  |  |  |
| --- | --- | --- | --- | --- | --- | --- | --- | --- | --- | --- | --- | --- | --- |
|  | P | R | F1 | P | R | F1 | P | R | F1 | P | R | F1 | support |
| eukaryote | 0.98 | 0.98 | 0.98 | 1.00 | 0.96 | 0.98 | 1.00 | 0.97 | 0.99 | 1.00 | 1.00 | 1.00 | 4411 |
| prokaryote | 0.94 | 0.95 | 0.95 | 0.88 | 1.00 | 0.94 | 0.99 | 1.00 | 1.00 | 1.00 | 1.00 | 1.00 | 1464 |
| organelle | - | - | - | - | - | - | 0.00 | 0.00 | 0.00 | - | - | - | 0 |
| unknown | - | - | - | - | - | - | 0.00 | 0.00 | 0.00 | - | - | - | 0 |
| Accuracy |  |  | 0.97 |  |  | 0.97 |  |  | 0.98 |  |  | 1.00 | 5875 |
| Macro avg | 0.96 | 0.97 | 0.97 | 0.94 | 0.98 | 0.96 | 0.50 | 0.49 | 0.50 | 1.00 | 1.00 | 1.00 | 5875 |
| Weighted avg | 0.97 | 0.97 | 0.97 | 0.97 | 0.97 | 0.97 | 1.00 | 0.98 | 0.99 | 1.00 | 1.00 | 1.00 | 5875 |

### Contig_classifier vs EukRep and Tiara

Because we wanted to make sure that our classifier (Contig_classifier) is also accurate on organisms that were not in our training set, we ran our classifier on another test dataset made from genomes of 10 eukaryotes and 15 prokaryotes (Supplementary Data 2), for which we prepared artificial contigs the same way as with the training dataset. This resulted in 4,411 eukaryotic contigs and 1,464 prokaryotic contigs. Most of the eukaryotic organisms in this dataset were also not used in the training data for EukRep and Tiara (Supplementary Data 2). The accuracy on this dataset was 97%, with a precision and recall of both 0.98 for eukaryotes and 0.94 and 0.95 for prokaryotes, respectively (Table 1). We also wanted to know how our classifier performs in comparison to EukRep and Tiara, which use k-mer counts of fragmented contigs as features for a support vector machine classifier and a neural network-based classifier, respectively. We ran our classifier, EukRep (default model, chunk size 5000 bp) and Tiara (default settings) on our test dataset to compare their performances. Both contig_classifier and EukRep had an overall accuracy of 97%, but our classifier had a precision of 0.94 for prokaryotic contigs, while EukRep had a much lower precision score of 0.88 (Table 1). Tiara had an overall accuracy score of 0.98, but a recall of 0.97 for eukaryotes (Table 1). It also identified sequences as ‘unknown’ or ‘organelle’, while we excluded organellar DNA from our dataset.

We then wanted to know if all three classifiers make similar mistakes. Firstly, we calculated the accuracy per bin of contig lengths with bin sizes of 10kbp. Contig_classifier and EukRep are less accurate on contigs shorter than 20kbp, but with at least 90% of contigs correctly classified, the accuracy is still relatively high (Supplementary Figure 2). Tiara has a more consistent performance across all contig lengths, although predictions on contigs shorter than 20kbp also tend to be slightly less accurate. For our classifier, the lower accuracy may stem from the fact that shorter contigs contain fewer genes based on which the features are calculated. The EukRep classifier may be less accurate on shorter contigs because these can be fragmented into fewer 5kbp pieces than longer contigs, making a majority vote to determine the overall contig class less reliable.

When we calculate the accuracy on contigs per organism, we see that each classifier has problems with different organisms, with no bias towards eukaryotes or prokaryotes (Figure 2). Both EukRep and Tiara make relatively many mistakes in eukaryotes, and fewer in prokaryotes (Figure 2). For example, EukRep only classifies about 75% of *Schizosaccharomyces pombe* contigs correctly, while all contigs of this organism were correctly classified by our classifier and Tiara. It is remarkable that EukRep performed relatively badly on *Schizosaccharomyces pombe*, because it was trained on this organism. The EukRep training genomes were not fragmented as much as those in our training set, however. If distinct k-mer frequencies are unevenly distributed across the genome, the distributions in larger contigs may be different than in shorter contigs. This may be a limitation of using k-mer counts as the only features. Tiara performed poorly on *Beta vulgaris*, a higher plant. This may be because Tiara did not train on higher plant nuclear genomes, but only on green algae. On the other hand, EukRep and Tiara performed better on *Xylella taiwanensis* contigs, with about 95% of contigs correctly classified, versus only ∼50% by our classifier (Figure 2). We expected that our model is less accurate in predicting the class of a contig when the genome structure of the organism is atypical and more resemblant of the other class. Indeed, about half of the contigs of *Xylella taiwanensis* have an intergenic distance and gene density that fall within a range that is closer to that of most eukaryotes (Supplementary Figure 3). We use a limited set of features that are all related, which means that if the gene structure in (part of) an organism’s genome is atypical, the classifier may not recognize this and assign the wrong class.

**Figure 2.**
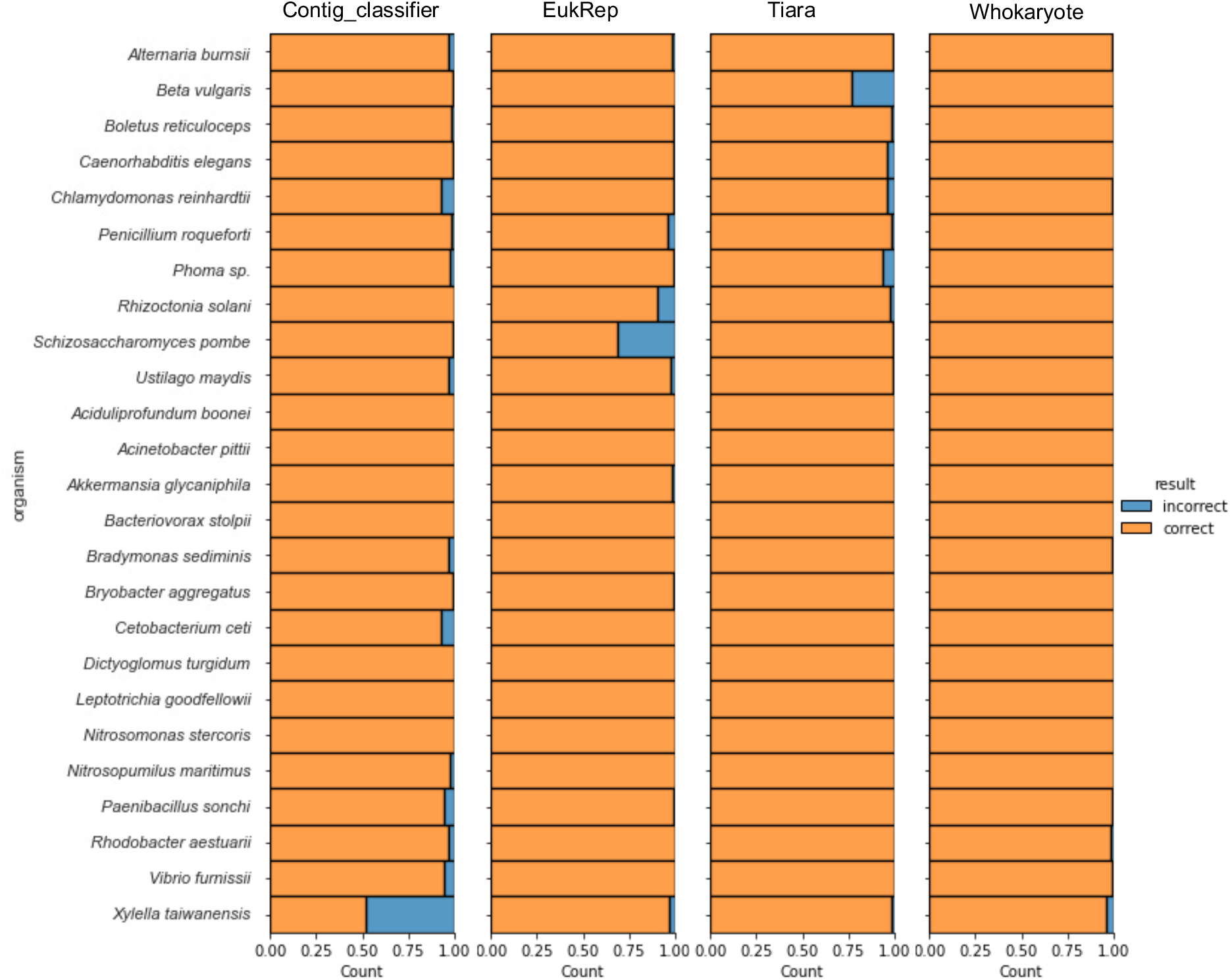
Comparison of the accuracy (depicted as relative count of incorrect and correct predictions) per organism of contig_classifier, EukRep, Tiara and Whokaryote on a test dataset with 10 eukaryotes and 15 prokaryotes.

### Enhanced classifier that uses tiara predictions as an additional feature is highly accurate

Because all three classifiers make mistakes in different contigs, it is possible that the important k-mers of EukRep and Tiara contain information complementary to the features based on gene structure that we have used. Therefore, we hypothesized that if we included the Tiara output as a feature to re-train our own classifier, the overall accuracy should improve. Indeed, retraining the classifier with the Tiara output from the first step, with ‘bacteria’ and ‘archaea’, and ‘unknown’ and ‘organelle’ collated into single classes, resulted in a classifier with an overall accuracy of 99.8%. Rounded F-1 scores are 1.00 for both eukaryotes and prokaryotes (Table 1).

The Tiara prediction was the most important single feature, with a score of 0.335, while gene density and intergenic distance in sum had a somewhat larger feature importance (with 0.2 and 0.19, respectively) (Supplementary Figure 4). The enhanced classifier improves the accuracy scores for the organisms that either of the individual classifiers performed weakly on, such as *Xylella fastidiosa* (Figure 2). This shows that by combining two classifiers with different feature types, the weaknesses of each classifier can be compensated, which results in a highly accurate new classifier. We call the resulting classifier ‘Whokaryote’.

The Tiara preprint reported a relatively high percentage of misclassifications in genomes from specific microbes with reduced genomes such as symbionts and parasites. We selected a set of such genomes to see if our initial classifier and Whokaryote would perform better. The genomes were divided into contigs of > 5000 bp, resulting in +-1600 contigs in total, varying between 8 and 495 contigs per organism ([Supplementary Data #]). On these contigs, we ran Tiara (standalone), our original contig_classifier, and Whokaryote (Supplementary Figure 5). The overall accuracy scores were 61% for Tiara, 80% for contig_classifier, and 86% for Whokaryote. Whokaryote performed better than Tiara on 7 out of 10 organisms, and similar on the other three organisms. Interestingly, our standalone initial classifier performed better than both Tiara and Whokaryote on 4 organisms. Tiara performed the worst on the eukaryotic microbe *Cafeteria roenbergensis* and on the bacterium *Mycoplasma haemofelis*, with < 25% of contigs classified correctly. Our initial ‘contig_classifier’ performed better on the contigs of these genomes, with +-70% and 55% of contigs classified correctly, respectively. Interestingly, Whokaryote performed better than contig_classifier and Tiara on *C. roenbergensis* (> 75% correct), but on *Mycoplasma haemofelis* artificial contigs it performed better than Tiara but worse than contig_classifier, with around 30% of contigs correctly classified. For the *Parcubacteria* metagenome-assembled genome, contig_classifier classified 100% of the artificial contigs correctly, while Tiara and Whokaryote both classified about 80% correctly. This indicates that in some cases, the addition of the Tiara prediction as a feature leads to worse performance of Whokaryote. When working with microbiomes that contain genomes of unusual (e.g., parasitic) organisms, it may be advisable to (also) use our standalone classifier.

### Comparison with a homology-based approach on a real-world dataset

To test if our enhanced classifier also works on real-world data, we ran it on a real metagenome dataset of the endophytic root microbiome of sugar beet. We ran metaProdigal on the metagenomic contigs > 5-kbp to use as input for our classifier, together with the DNA sequences of the contigs for the Tiara classification. A total of 29,512 contigs with more than 1 gene were classified using a single core. The Tiara classification step took 226.7 seconds, and the rest of the classification took 95.2 seconds, resulting in a total time of 321.9 seconds (approx. 5 minutes). The classifier predicted 1,653 eukaryotic contigs and 27,859 prokaryotic contigs. We wanted to know if these classifications were reliable and compared them to the classification of homology-based classification tool CAT^10^ (standard settings, -n 20, nr database version 2021-01-07), which can taxonomically classify contigs up to the species level, but which took a much longer time, approximately 6 hours and 49 minutes, to run (excluding gene prediction) on 20 cores. However, we only looked at the classifications on the superkingdom level, and we labelled the classifications “Bacteria” and “Archaea” as ‘prokaryote’, and “Eukaryota” as ‘eukaryote’. Out of 29,512 contigs, 28,781 (97.5%) got classified with the same taxonomic label by both our classifier and CAT. Of the 731 contigs that did not have a matching classification, 163 were classified as ‘not_assigned’ by CAT, 351 were classified as ‘no support’ and 6 contigs were classified as Viruses. When CAT cannot find a match for an ORF in a database, it cannot assign any taxonomic information to the contig, and therefore classifies it as ‘no support’ or ‘not_assigned’. With our classifier, we were able to classify these contigs as having a prokaryotic or eukaryotic origin. Of the remaining non-matching classifications, CAT classified 146 contigs as prokaryote while our classifier identified them as ‘eukaryote’, and 65 contigs were classified as eukaryote by CAT and as prokaryote by our classifier. All in all, these results show that our enhanced classifier is also very accurate on real metagenomic data, and can quickly determine the portion of eukaryotic sequences, which can then be processed with eukaryote-specific tools.

### More fungal biosynthetic gene clusters detected on fungal contigs with re-predicted genes

EukRep already showed that binning of eukaryotic contigs helps improve gene predictions and functional annotations. However, there will always be contigs that cannot be binned. These may still contain interesting genes that may help explain the functions of the microbiome. The secondary metabolites/natural products that are produced by microbes are of special interest in this regard, as these molecules are often used to interact with other microbes and the environment. The biosynthetic gene clusters (BGCs) encoding the production of these molecules can be predicted by tools such as antiSMASH^20^, which uses specific parameters and models for predicting bacterial BGCs, and a different set of parameters and models to predict fungal BGCs. Additionally, the bacterial version uses Prodigal as a gene finder, and the fungal version (fungiSMASH) uses GlimmerHMM^21^, an ab initio eukaryotic gene finder. Currently, many metagenomic studies do not distinguish between fungal and bacterial contigs and run the bacterial version of antiSMASH on all contigs. Important fungal BGCs may be missed because of wrong gene annotations and the use of bacteria-specific models. We wanted to know if pre-filtering metagenomic contigs into eukaryotic and prokaryotic could be a useful approach to select contigs that should (also) be run using the fungal mode of antiSMASH. We ran both the bacterial version (options --genefinding-tool prodigal-m) and the fungal version (options -- taxon fungi, --cassis, --genefinding tool glimmerhmm) of antiSMASH on the contigs of our test dataset and compared the results. The bacterial version found 120 regions, of which 65 were located on eukaryotic contigs and 55 on prokaryotic contigs. The fungal version found 86 regions, of which 70 were located on eukaryotic contigs and 16 on prokaryotic contigs. 20 eukaryotic BGCs were found by fungiSMASH that were not found by antiSMASH, showing that using eukaryote-specific tools on contigs classified as eukaryotic can lead to the discovery of functions that are missed when using prokaryote-specific pipelines to study metagenomes. Interestingly, bacterial antiSMASH also found BGCs on 15 eukaryotic contigs that were not found by fungiSMASH.

We also ran antiSMASH and fungiSMASH (version 5.1.2) on the sugar beet endophyte metagenome. AntiSMASH found 31 BGCs on eukaryotic contigs, while fungiSMASH found 26 BGCs on eukaryotic contigs. FungiSMASH reported 4 BGCs on eukaryotic contigs that were not found by antiSMASH, namely a type-one polyketide synthase (T1PKS), a terpene, an NRPS-like, and a dual cluster with non-ribosomally translated peptide synthase-like and a terpene (Figure 3). At the same time, there were also 9 BGCs on eukaryotic contigs that were only predicted by antiSMASH and not by fungiSMASH. This shows that simply using a general eukaryote-specific gene predictor (GlimmerHMM) that was trained on a single species, as implemented in antiSMASH, on eukaryotic contigs does not necessarily lead to better gene predictions for BGC detection. BGC core genes, which are scanned for signature domains, are usually very large and may contain multiple domains. Possibly, these genes are more difficult to predict correctly for eukaryotic gene finders. Nevertheless, some new BGCs were found that would not have been found with bacterial antiSMASH. Therefore, we suggest using both Prodigal and eukaryotic gene predictors (e.g., GlimmerHMM^21^, Augustus^13^, MetaEuk^15^) to predict genes on standalone eukaryotic contigs and use both annotations for further functional analyses.

**Figure 3.**
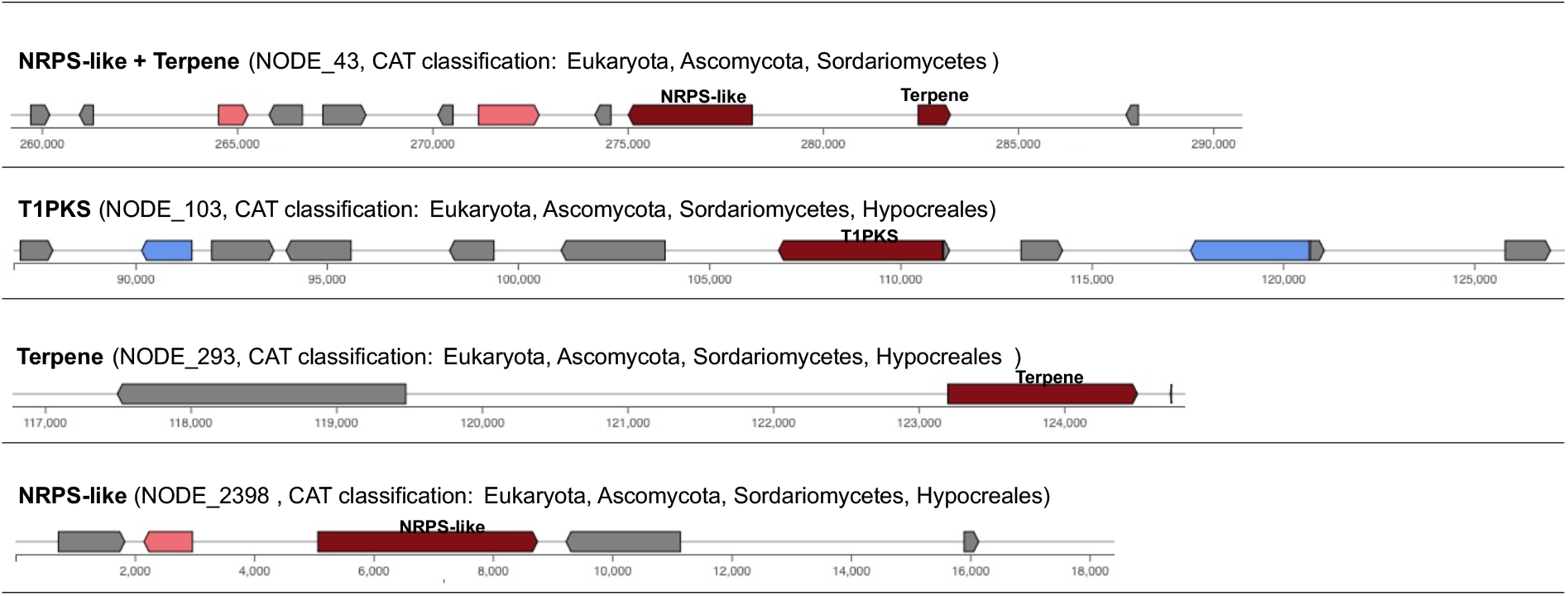
Four BGCs on beet endophyte metagenome contigs classified as eukaryotic by Whokaryote. These four BGCs were predicted by fungiSMASH, which uses GlimmerHMM as a gene predictor, but were not found with antiSMASH, which uses Prodigal-M as a gene predictor.

## Discussion

Studying eukaryotes in metagenomes is still challenging, and tools that aid with this process are scarce and not widely used yet. First, we made a random forest classifier that reliably determines whether a metagenomic contig (or any genomic sequence) is of eukaryotic or prokaryotic origin by using features based on fundamental differences between the genome structures of prokaryotes and eukaryotes. Features are calculated with Prodigal genome annotation files. Prodigal is not made for predicting eukaryotic genes and does not detect introns and may therefore predict multiple exons of a single gene as multiple smaller complete genes. While Prodigal annotations are suited for our classification goal, it is important to keep in mind that the classifier is based on differences in features that arise partly from inadequately annotating eukaryotic contigs. Nevertheless, the differences in features we see between eukaryotes and prokaryotes (except for gene length) are in line with current knowledge of the gene structures of these organisms, and the importance of those features to our classifier support this. The classifier struggled with contigs from a prokaryote that had genomic regions more resembling the gene structure of eukaryotes.

We compared our classifier to EukRep and Tiara, which use k-mer counts with a support vector machine or a neural network, respectively, to classify contigs on the eukaryote/prokaryote level. In general, Tiara outperforms both our initial classifier and EukRep, but when looking at the results per organism we can see that Tiara and EukRep more often misclassify specific eukaryotes, while our initial classifier has more misclassifications in prokaryotes. Additionally, all three classifiers perform worse on shorter contigs, but with an average accuracy of over 90% on contigs <10 kb the results can still be used on a large number of contigs in high-quality metagenome assemblies. For both EukRep and Tiara, it is unknown which k-mers are the most important for classification and if they correspond to any known biological signal. Our classifier makes use of manually selected features based on prior biological knowledge and on its own, performs (almost) equally well as the other classifiers. However, all three tools make different mistakes, indicating that the k-mers represent sequence motifs that do not necessarily correspond to our features. We speculate that gene-structure-based features may be more general because of the fundamental biological differences between eukaryotes and prokaryotes. Methods that use k-mer frequencies may pick up on signals, such as specific sequence motifs, that are more specific to higher taxonomic ranks and may be more accurate on genomes that are closely related to training set sequences. A very wide training database, such as those used by Tiara and EukRep, may alleviate this problem. Our relatively small training dataset and the positive results on our test dataset confirm that our features are also predictive for genomes more distantly related than those of the training set.

By re-training our initial classifier with Tiara predictions as additional feature, weaknesses of both classifiers are compensated, and the result is an enhanced classifier that almost flawlessly classifies contigs as eukaryotic or prokaryotic on our test dataset, while still being very fast. Additionally, on specific challenging genomes, our initial classifier and Whokaryote show improved accuracy when compared to Tiara. On a real-world metagenome of a plant root endophytic microbiome, Whokaryote is very accurate, and it can classify contigs of organisms that are not in any database yet and can thus not be classified via sequence homology. Although the enhanced classifier uses two different classifiers, it is still very fast with just over 5 minutes to run on a single core on a metagenome consisting of 29,512 contigs and totaling 0.49 Gbp. In comparison, CAT took almost 7 hours to align and taxonomically classify the contigs on 20 cores.

Previous findings showed that using a eukaryotic gene predictor on a metagenome-recovered eukaryotic bin resulted in a more complete predicted gene set when compared to MetaProdigal predictions^14^. We show that re-predicting genes on unbinned contigs is also worthwhile and can lead to the discovery of genes and functions that would not have been found otherwise. Future research should focus on more accurate ab initio eukaryotic gene prediction on unbinned contigs to prevent these extra steps.

## Conclusions

We show that using manually selected features based on fundamental differences in gene structure between eukaryotes and prokaryotes can be used to reliably classify metagenomic contigs as eukaryotic or prokaryotic. Using only these features leads to similar accuracies as k-mer frequency-based approaches. With Whokaryote, we combined our selected features with the output from the k-mer based deep learning classifier Tiara. The resulting classifier achieves nearly flawless classification of contigs from a wide range of organisms and compensates biases present in the individual classifiers. Contigs predicted as eukaryotic can be included in metagenomic pipelines by using eukaryote-specific tools, allowing new discoveries about their roles in microbiomes.

## Supplementary Figures

**Supplementary Figure 1.**
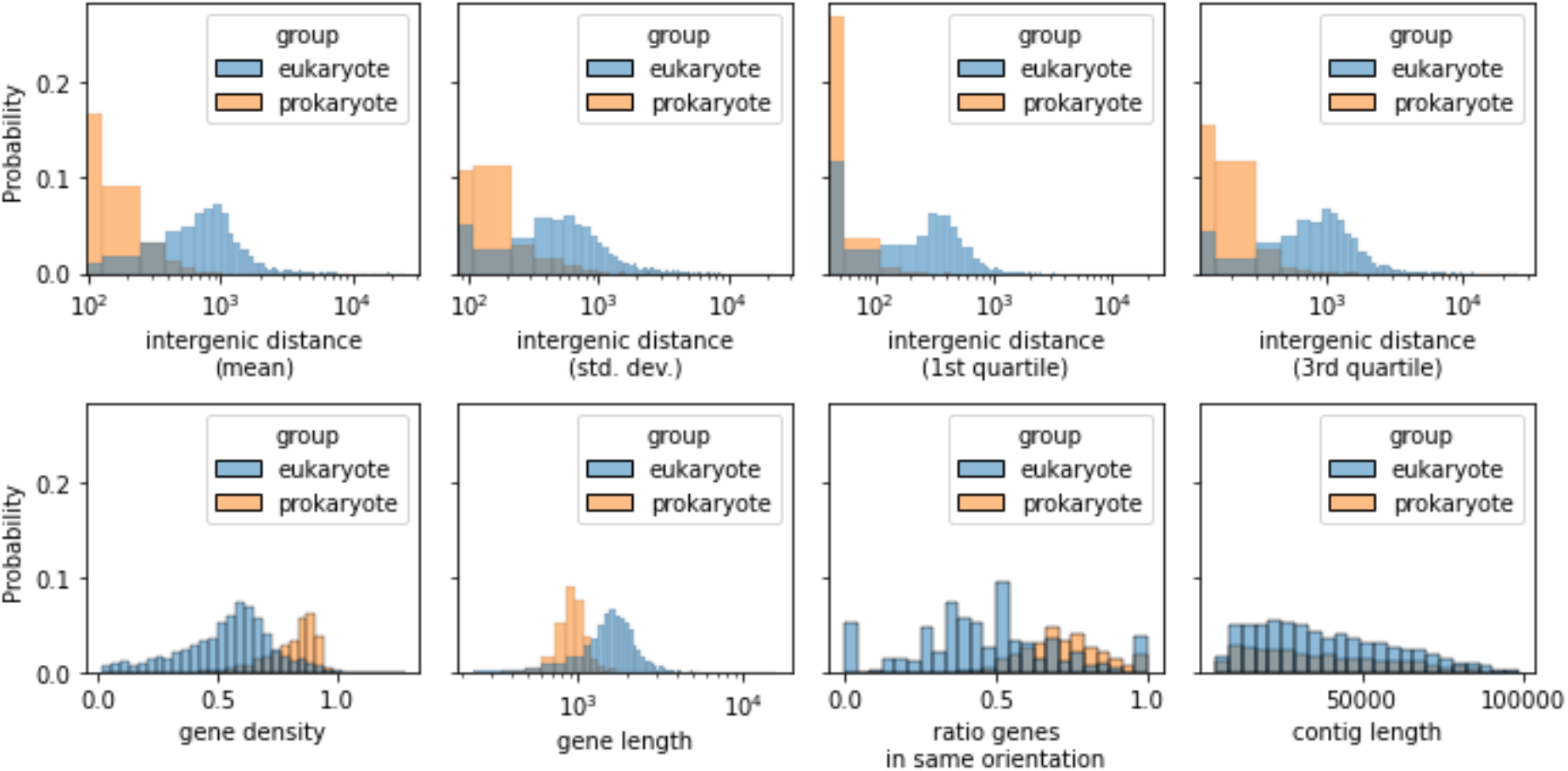
Overview statistics of the features calculated for the training and validation dataset containing artificial contigs taken from 73 prokaryotic and 25 eukaryotic reference genomes with their reference gene annotations. The probability distribution of the feature values of all contigs are shown, grouped by the taxonomic group (eukaryote or prokaryote) the contig belongs to. For every contig, the mean, standard deviation, first quartile and third quartile of the intergenic distances between the genes was calculated (a-d, respectively). e: The mean gene density of every contig, calculated by dividing the sum of the length (in basepairs) of all genes by the total length of the contig. f: The mean gene length of every contig, calculated as the end position of the gene minus the start position. g: The ratio of genes on a contig that are on the same strand (e.g., they have the same ‘orientation’).

**Supplementary Figure 2.**
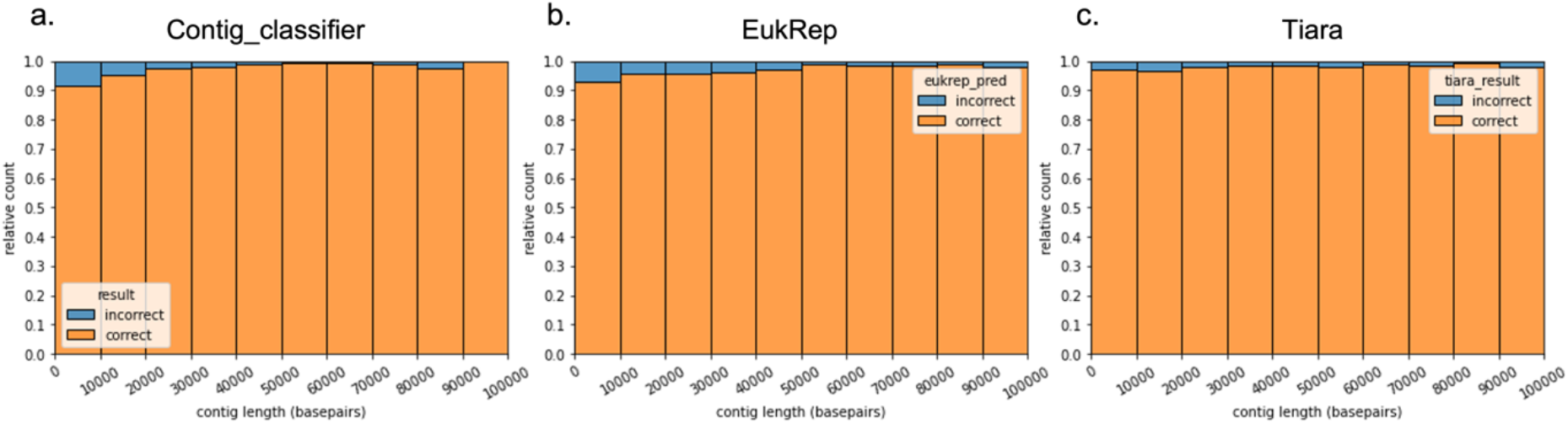
The classification accuracy in different contig length categories of contig_classifier, EukRep and Tiara, shown as relative count of incorrect and correct predictions.

**Supplementary Figure 3.**
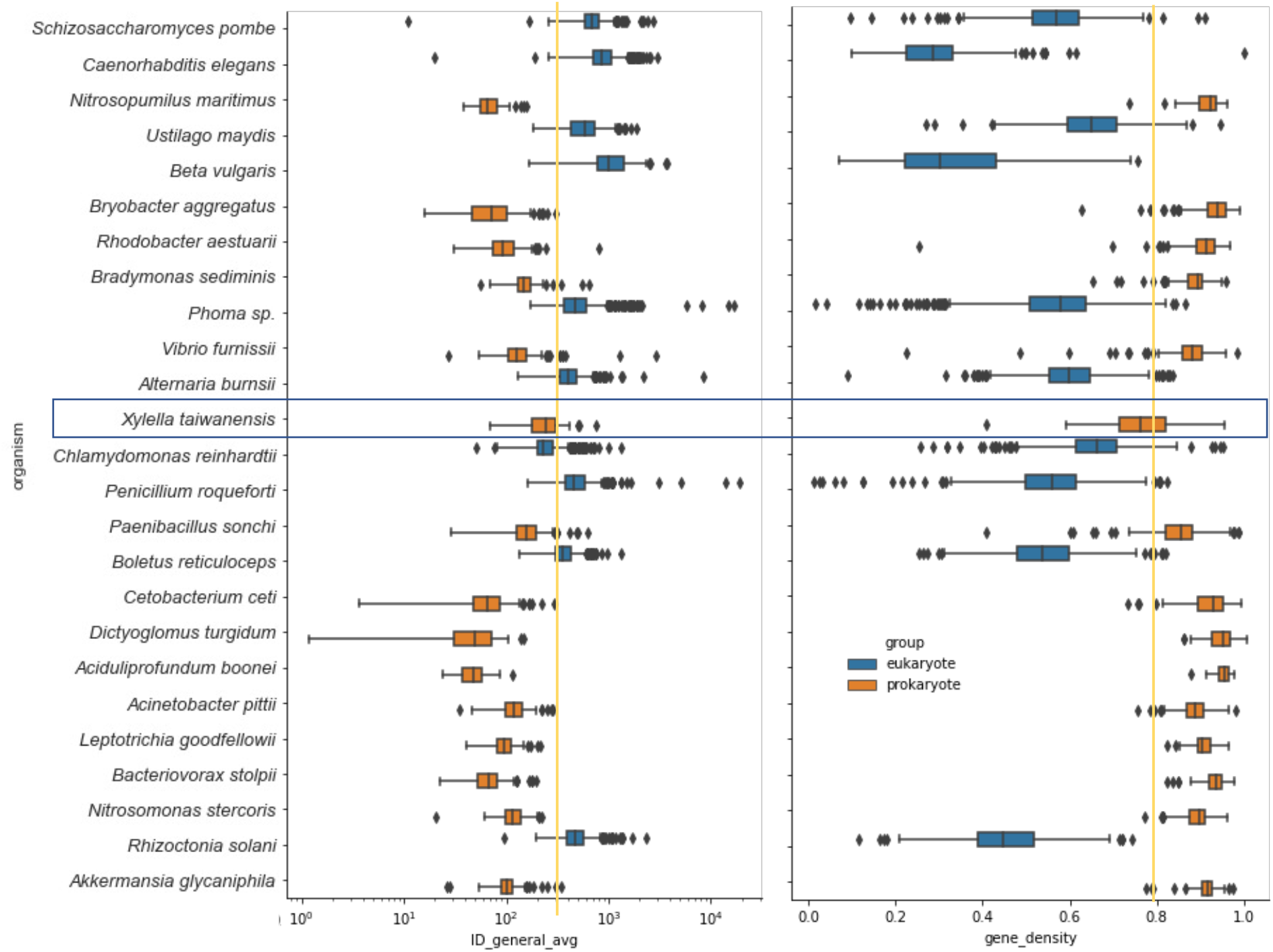
On the left, the distribution of average intergenic distances per contig is shown per organism. A vertical yellow line was drawn roughly between the eukaryotic and prokaryotic boxes. A blue box is drawn around the results for Xylella taiwanensis to show that the average intergenic distance of its contigs lies very close to the range of most eukaryotes. On the right, the same is shown for the average gene density.

**Supplementary Figure 4.**
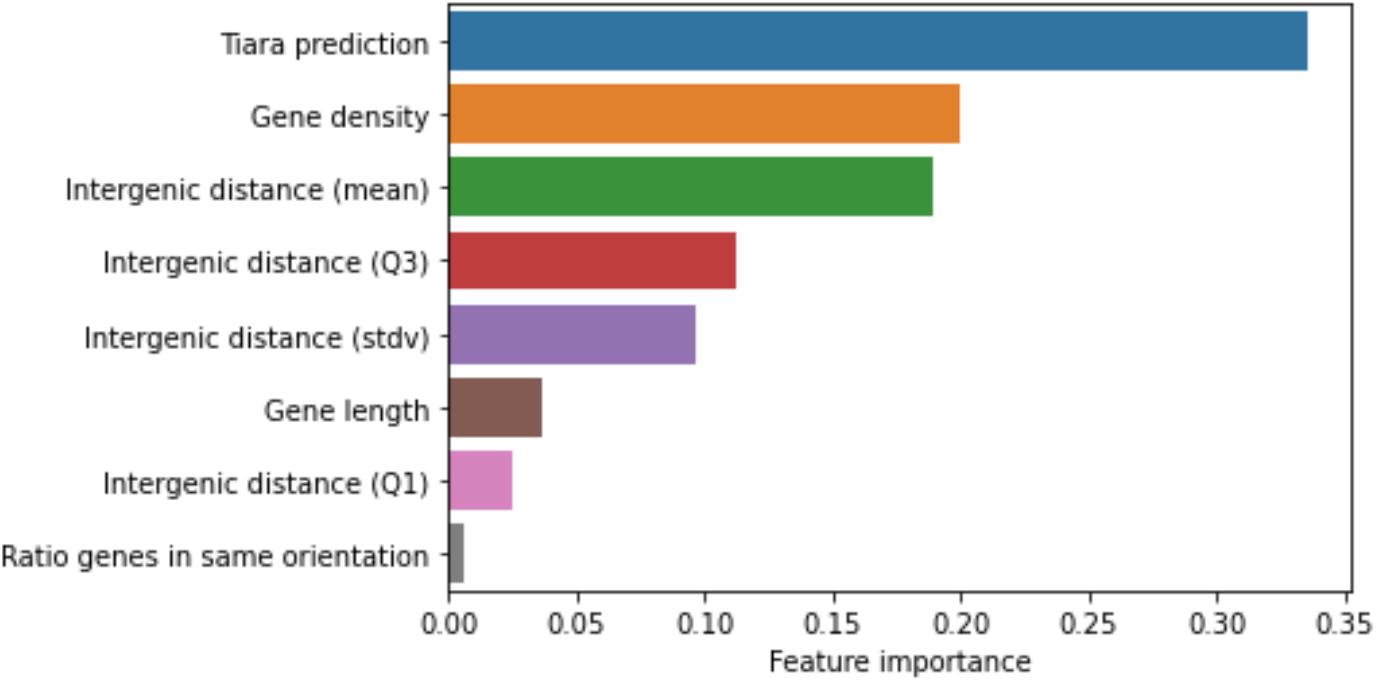
Feature importance of the enhanced random forest classifier ‘Whokaryote’, which uses Tiara predictions (eukaryote/prokaryote) as an extra feature.

**Supplementary Figure 5.**
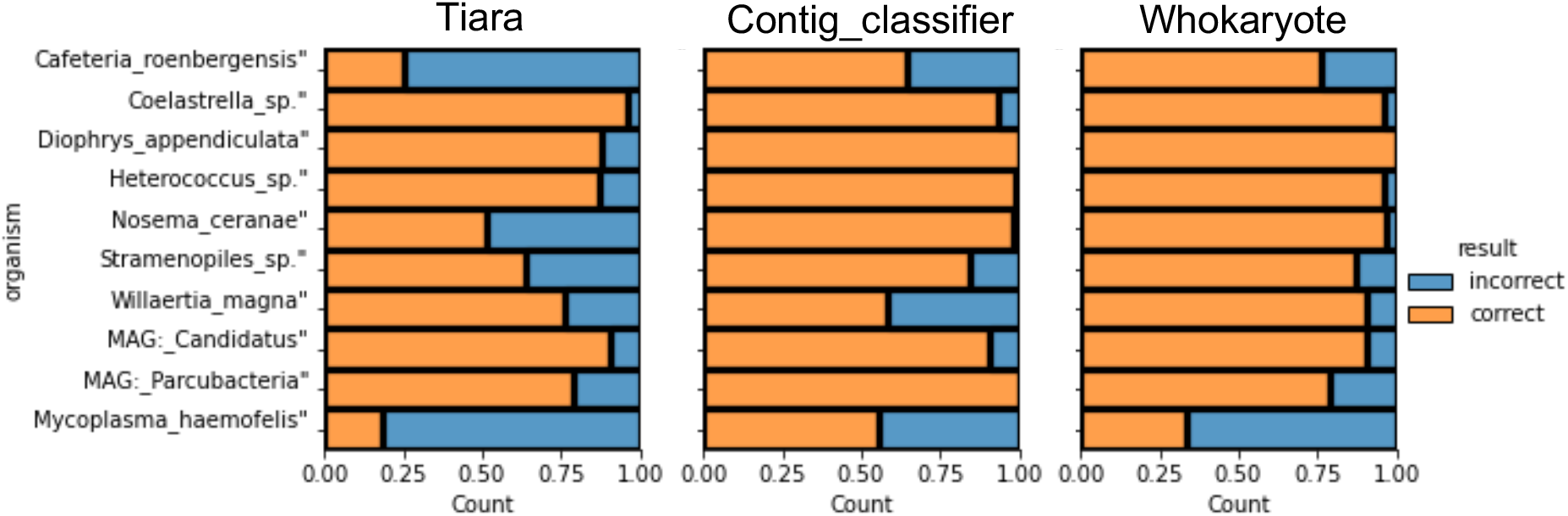
Tiara, contig_classifier and Whokaryote were tested on a set of genomes on which Tiara performed relatively badly. The genomes of these organisms were split into random contigs between 5000 and 100000 basepairs, with a total of around 1600 contigs.

## Notes

### Competing Interest Statement

M. H. M. is a co-founder of Design Pharmaceuticals and a member of the scientific advisory board of Hexagon Bio.

